# *Olsenella lakotia* SW165 sp. nov., an acetate producing obligate anaerobe with a GC rich genome

**DOI:** 10.1101/670927

**Authors:** Supapit Wongkuna, Sudeep Ghimire, Roshan Kumar, Linto Antony, Surang Chankhamhaengdecha, Tavan Janvilisri, Joy Scaria

## Abstract

A Gram-positive and obligately anaerobic bacterium was isolated from cecal content of feral chickens in Brookings, South Dakota, USA. The microorganism grew at 37-45° C and pH 6-7.5. This strain produced acetic acid as the primary metabolic end product. Major fatty acids were C_12:0_, C_14:0_, C_14:0_ DMA and summed feature 1 (C_13:1_ at 12-13 and C_14:0_ aldehyde). Phylogenetic analyses based on 16S rRNA gene sequence suggested that strain SW165 belongs to the family *Atopobiaceae* with the closest relatives being *Olsenella profusa* DSM 13989^T^ (96.33% similarity), *Olsenella umbonate* DSM 26220^T^ (96.18%) and *Olsenella uli* DSM 7084^T^ (96.03%). Genome sequencing revealed a genome size of 2.43 Mbp with a G+C content of 67.59 mol%, which is the highest G+C content among members of the genus *Olsenella*. Phylogenetic and phenotypic comparison indicated that strain SW165 represents a novel species of the genus *Olsenella*, for which the name *Olsenella lakotia* sp. nov. is proposed. The type strain is SW165 (=DSM 107283^T^).

## Introduction

The members of the genus *Olsenella* are strictly anaerobic, gram-stain-positive, non-motile, non-spore-forming rods or cocci and was first named by Dewhirst *et al.* 2001 (1). This genus has recently been reclassified as a member of the family *Atopobiaceae*, order *Coriobacteriales*, class *Coriobacteriia*, and phylum *Actinobacteria* (2). The genus *Olsenella* consists of nine species; *Olsenella uli* (Olsen *et al.* 1991) (3), *Olsenella profusa* (Dewhirst *et al.* 2001) (1), *Olsenella umbanata* (Kraatz *et al.* 2011) (4), *Olsenella scatoligenes* (Li *et al.* 2015) (5), *Olsenella urininfantis* (Morand et al. 2016) (6), *Olsenella congonensis* (Bilen et al. 2017) (7), *Olsenella provencensis, Olsenella phocaeensis*, and *Olsenella mediterranea* (Ricaboni et al. 2017) (8). The main habitats of the *Olsenellae* are the oral cavity and gastrointestinal tract of humans (9-12), animals and various anaerobic environmental sites (13-15).

The chicken intestine harbors diverse microbial species. Many members of genus *Olsenellae* have been reported in the chicken microbiome in metagenomic-based studies (7, 16). However, only *Olsenella uli* was isolated from the chicken gut (17). Here, we describe strain SW165, which was isolated from the cecum of feral chicken and identified it as new species for which name *Olsenella lakotia* sp. nov. is proposed.

## Isolation and Ecology

Strain SW165 was isolated from the cecum of feral chicken. For cultivation, fresh cecal content was transferred within 10 minutes of the collection into an anaerobic workstation (Coy Laboratory) containing 85% nitrogen, 10% hydrogen and 5□% carbon dioxide. Modified Brain Heart Infusion (BHI-M) medium was used for strain isolation containing 37 g/L of BHI, 5 g/L of yeast extract, 1 ml of 1 mg/mL menadione, 0.3 g of L-cysteine, 1 mL of 0.25 mg/L of resazurin, 1 mL of 0.5 mg/mL hemin, 10 mL of vitamin and mineral mixture, 1.7 mL of 30 mM acetic acid, 2 mL of 8 mM propionic acid, 2 mL of 4 mM butyric acid, 100 µl of 1 mM isovaleric acid, and 1% of pectin and inulin. The strain was maintained in BHI-M medium and stored with 10% (v/v) Dimethyl Sulfoxide (DMSO) at −80° C. Reference strain *Olsonella profusa* DSM 13989^T^ was obtained from the German collection of microorganisms and cell culture was maintained under the same conditions.

## Physiology and Chemotaxonomy

For morphological characterization, the strain was anaerobically cultivated in BHI-M medium, pH 6.8-7.2, at 37° C. Colony morphologies were determined after 2-3 days of incubation on BHI-M agar plates. Gram-staining was performed using a Gram-Straining kit set (Difco), according to the manufacturer’s instructions. Cell morphologies were examined by scanning electron microscopy (SEM) of cultures during exponential growth. Aerotolerance was examined by incubating cultures for 2 days separately under aerobic and anaerobic conditions. Growth of strain SW165 at 4, 20, 30, 37, 40 and 55° C was determined. For determining the pH range for growth, the pH of the medium was adjusted to pH□4–9 with sterile anaerobic stock solutions of 0.1□M HCl and 0.1M NaOH. Motility of this microorganism was determined using motility medium with triphenyltetrazolium chloride (TTC) (29). The growth was indicated by the presence of red color, reduced form of TTC after it is absorbed into the bacterial cell wall.

Morphologically, strain SW165 matched descriptions of species of the genus *Olsenella*. Cells were gram-stain-positive bacilli (0.5–2.0 µm), growing in pairs or as short chains and were non-motile (Fig. 1 and Table 1). Colonies on BHI-M agar were 0.2-0.5 cm in diameter, appear white, smooth, and umbonate with entire circular edges. Strain SW165 grew between 37° C and 45° C with optimum growth at 45° C. The optimum pH for the growth was 7, and growth was observed at pH□6–7.

**Table 1.**
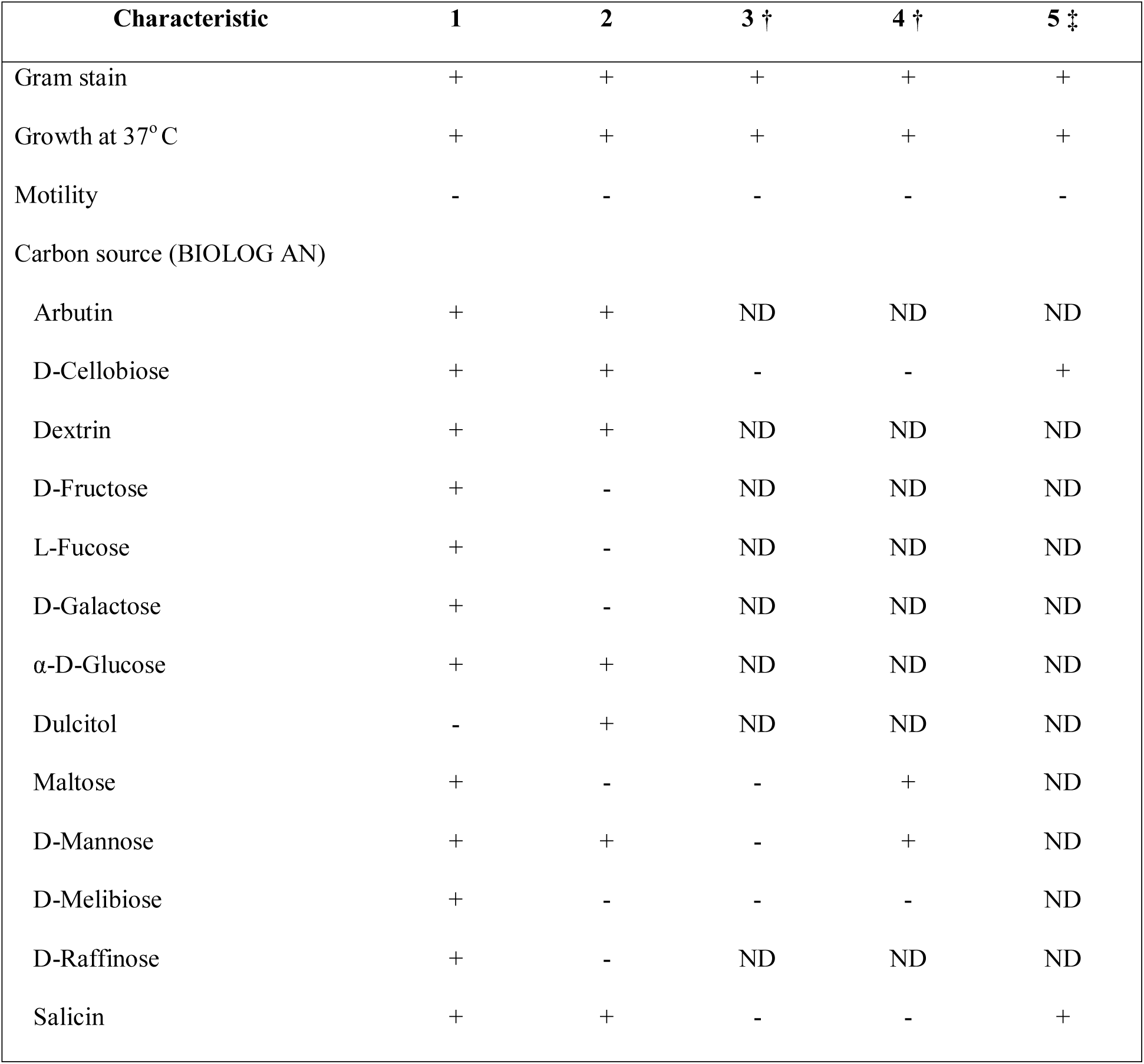
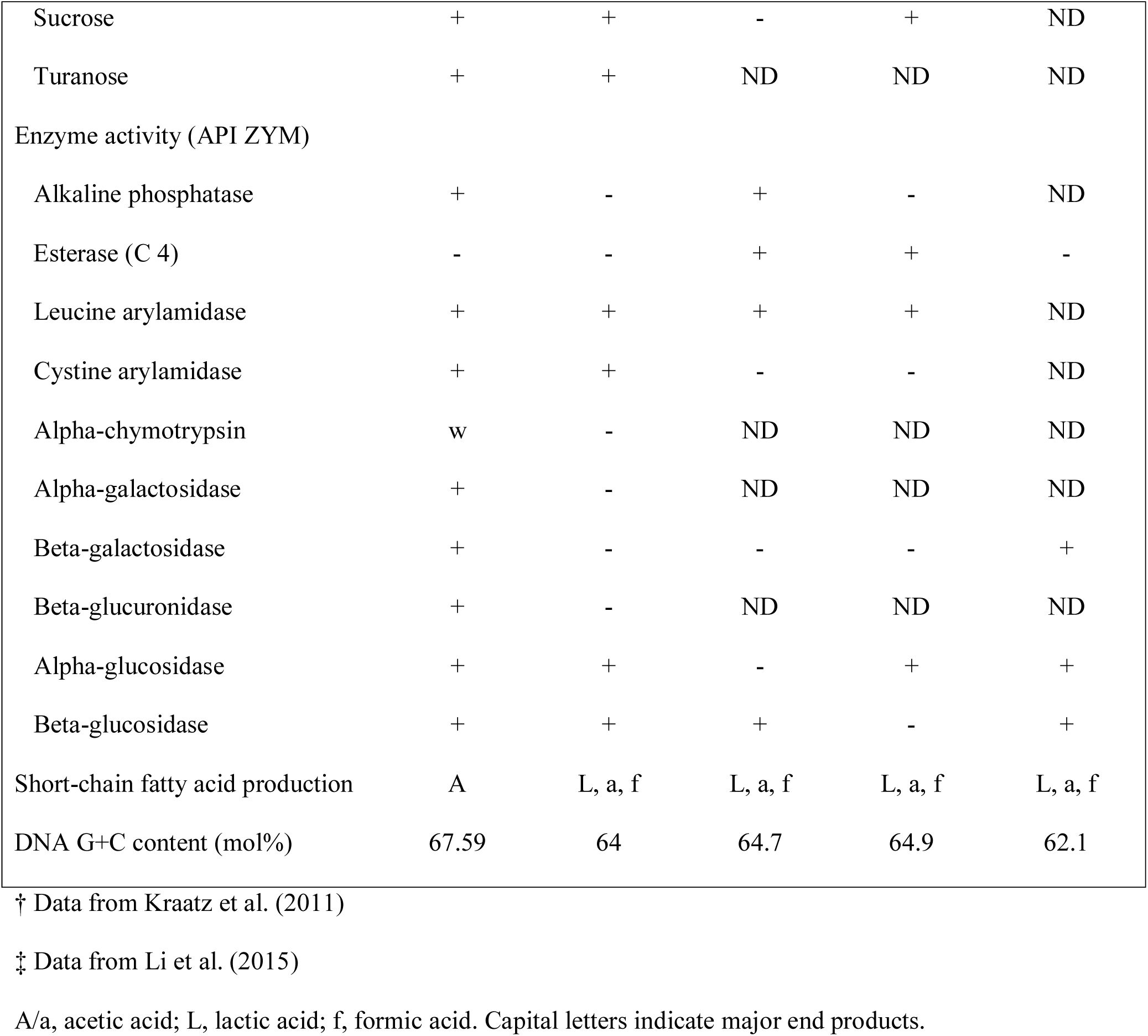
Characteristics of SW165 and closely related strains Column headers show Strains designated in the following numbers: 1 – SW165; 2 – *O. profusa* DSM 13989^T^; 3 – *O. uli* DSM 7084^T^; 4 – *O. umbanata* DSM 22620^T^; 5 – *O. scatoligenes* DSM 28304^T^. Results for metabolic end products of SW165 are from this study with cells that were cultured for 3 days at 37° C in BHI-M. +, positive; -, negative; w, weak; ND, not determined.

| Characteristic | 1 | 2 | 3 † | 4 † | 5 ‡ |
| --- | --- | --- | --- | --- | --- |
| Gram stain | + | + | + | + | + |
| Growth at 37° C | + | + | + | + | + |
| Motility | - | - | - | - | - |
| Carbon source (BIOLOG AN) |  |  |  |  |  |
| Arbutin | + | + | ND | ND | ND |
| D-Cellobiose | + | + | - | - | + |
| Dextrin | + | + | ND | ND | ND |
| D-Fructose | + | - | ND | ND | ND |
| L-Fucose | + | - | ND | ND | ND |
| D-Galactose | + | - | ND | ND | ND |
| α-D-Glucose | + | + | ND | ND | ND |
| Dulcitol | - | + | ND | ND | ND |
| Maltose | + | - | - | + | ND |
| D-Mannose | + | + | - | + | ND |
| D-Melibiose | + | - | - | - | ND |
| D-Raffinose | + | - | ND | ND | ND |
| Salicin | + | + | - | - | + |
| Sucrose | + | + | - | + | ND |
| Turanose | + | + | ND | ND | ND |
| Enzyme activity (API ZYM) |  |  |  |  |  |
| Alkaline phosphatase | + | - | + | - | ND |
| Esterase (C 4) | - | - | + | + | - |
| Leucine arylamidase | + | + | + | + | ND |
| Cystine arylamidase | + | + | - | - | ND |
| Alpha-chymotrypsin | w | - | ND | ND | ND |
| Alpha-galactosidase | + | - | ND | ND | ND |
| Beta-galactosidase | + | - | - | - | + |
| Beta-glucuronidase | + | - | ND | ND | ND |
| Alpha-glucosidase | + | + | - | + | + |
| Beta-glucosidase | + | + | + | - | + |
| Short-chain fatty acid production | A | L, a, f | L, a, f | L, a, f | L, a, f |
| DNA G+C content (mol%) | 67.59 | 64 | 64.7 | 64.9 | 62.1 |

† Data from Kraatz et al. (2011)
‡ Data from Li et al. (2015)
A/a, acetic acid; L, lactic acid; f, formic acid. Capital letters indicate major end products.

**Fig. 1.**
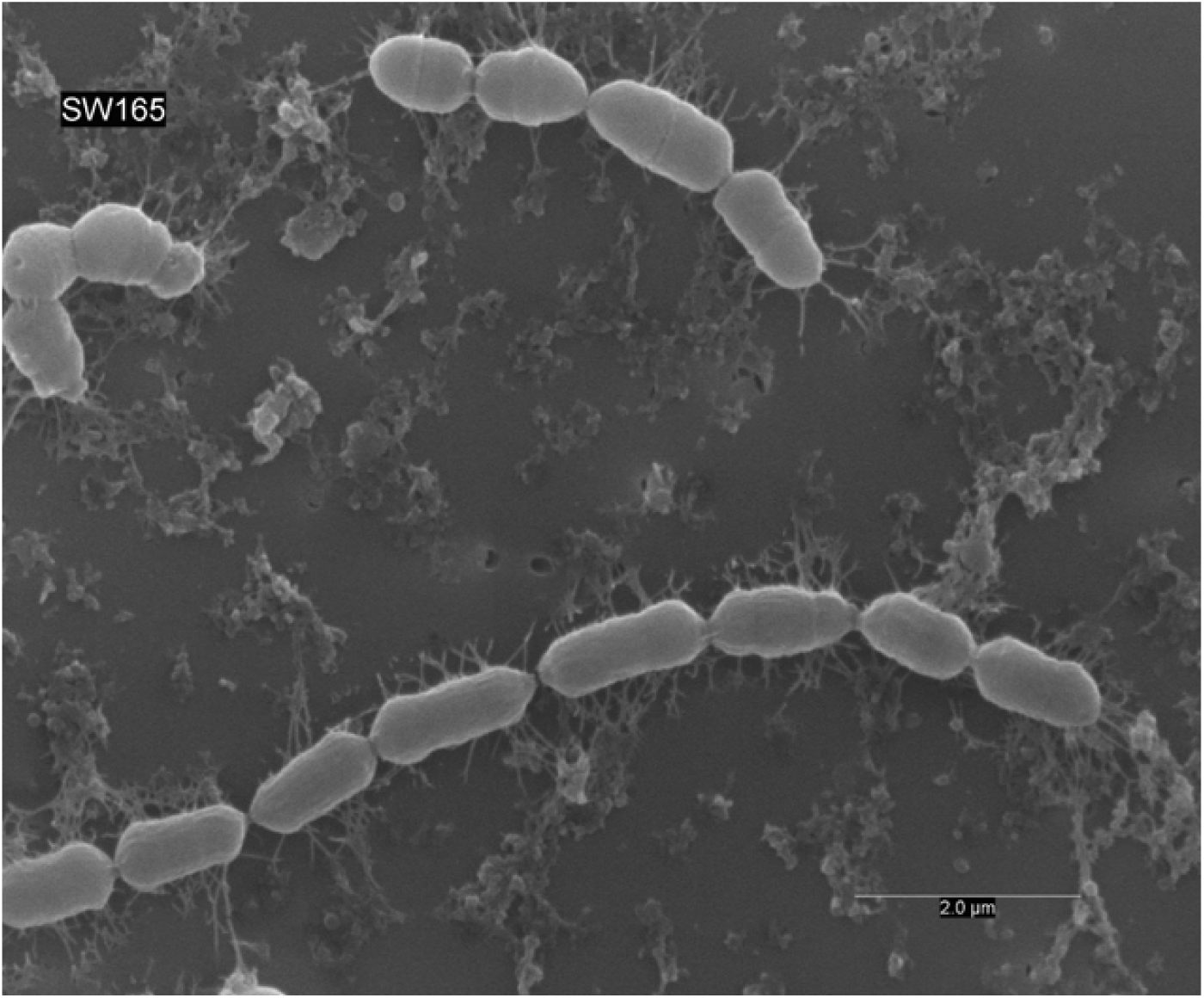
Scanning electron micrograph of strain SW165. Cells were anaerobically cultured for 24 hours at 37° C in BHI-M medium. Bar, 2 μm

The strain could grow well in BHI-M broth and on BHI-M agar under anaerobic conditions but not under aerobic conditions. This confirmed that the strains were obligately anaerobic. Biochemical tests to determine standard taxonomic characteristics were performed in triplicate. The utilization of various substrates as sole carbon and energy sources and enzyme activities were performed using the AN MicroPlate (Biolog) and API ZYM (bioMérieux) according to the manufacturer’s instructions. Strain SW165 and DSM 13989^T^ were simultaneously cultured in BHI-M medium at 37° C for 24□h under anaerobic condition before cell biomass was harvested for cellular fatty acids. The fatty acids were extracted, purified, methylated, identified and analyzed by GC (Agilent 7890A) according to the manufacturer’s instruction of Microbial Identification System (MIDI) (30). Metabolic end-products such as short-chain fatty acids of strain SW165 and DSM 13989^T^ grown in BHI-M were determined using gas chromatography. The cultures were maintained in 25% metaphosphoric acid before supernatant collection for the analysis. These analyses detected acetic acid, butyric acid, isovaleric acid and propionic acid.

Based on the results obtained in Biolog tests, strain SW165 differs from its closest phylogenetic neighbor in the utilization of D-fructose, L-fucose, D-galactose, maltose, D-melibiose and D-raffinose, and in the non-utilization of dulcitol. The strain SW165 exhibited positive detection of alkaline phosphatase, leucine arylamidase, cysteine arylamidase, α-galactosidase, β-galactosidase, β-glucuronidase, α-glucosidase and β-glucosidase, distinguishing from the closest strain (Table 1). The predominant cellular fatty acids of strain SW165 included C _12□:□0_ (25.5%), C _14□:□0_ (22.83%), C _14□:□0_ DMA (15.61%) and summed feature 1 [C_13:1_ and/or C_14:0_ aldehyde; 13.94%]. However, there were distinct quantities of some fatty acids between SW165 and the reference strain (Table 2). SW165 cultivated in BHI-M produced acetic acid (3.74 mM) and propionic acid (0.53 mM). Thus, the major volatile fatty acid produced by the strain SW165 was acetic acid.

**Table 2.**
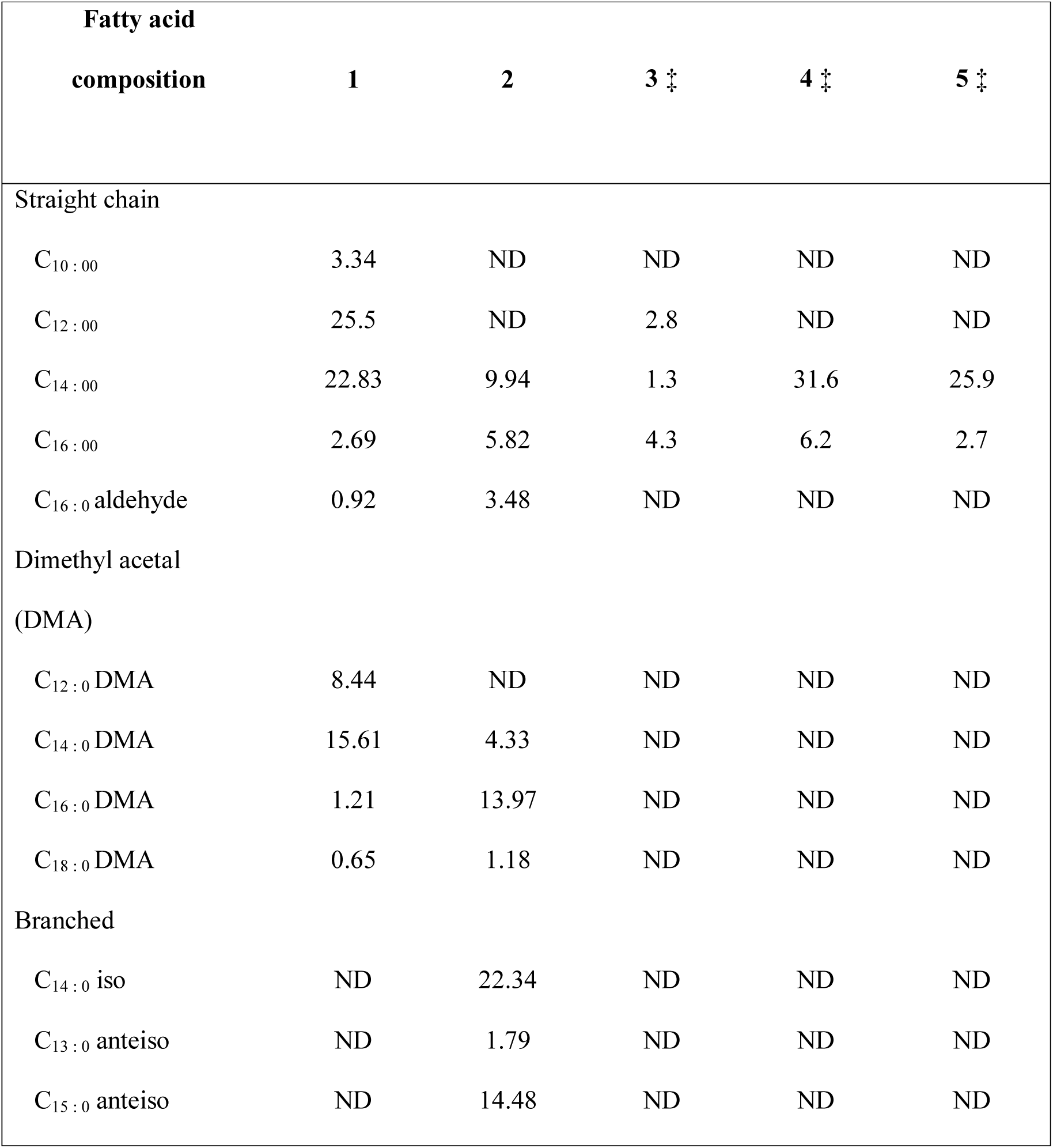
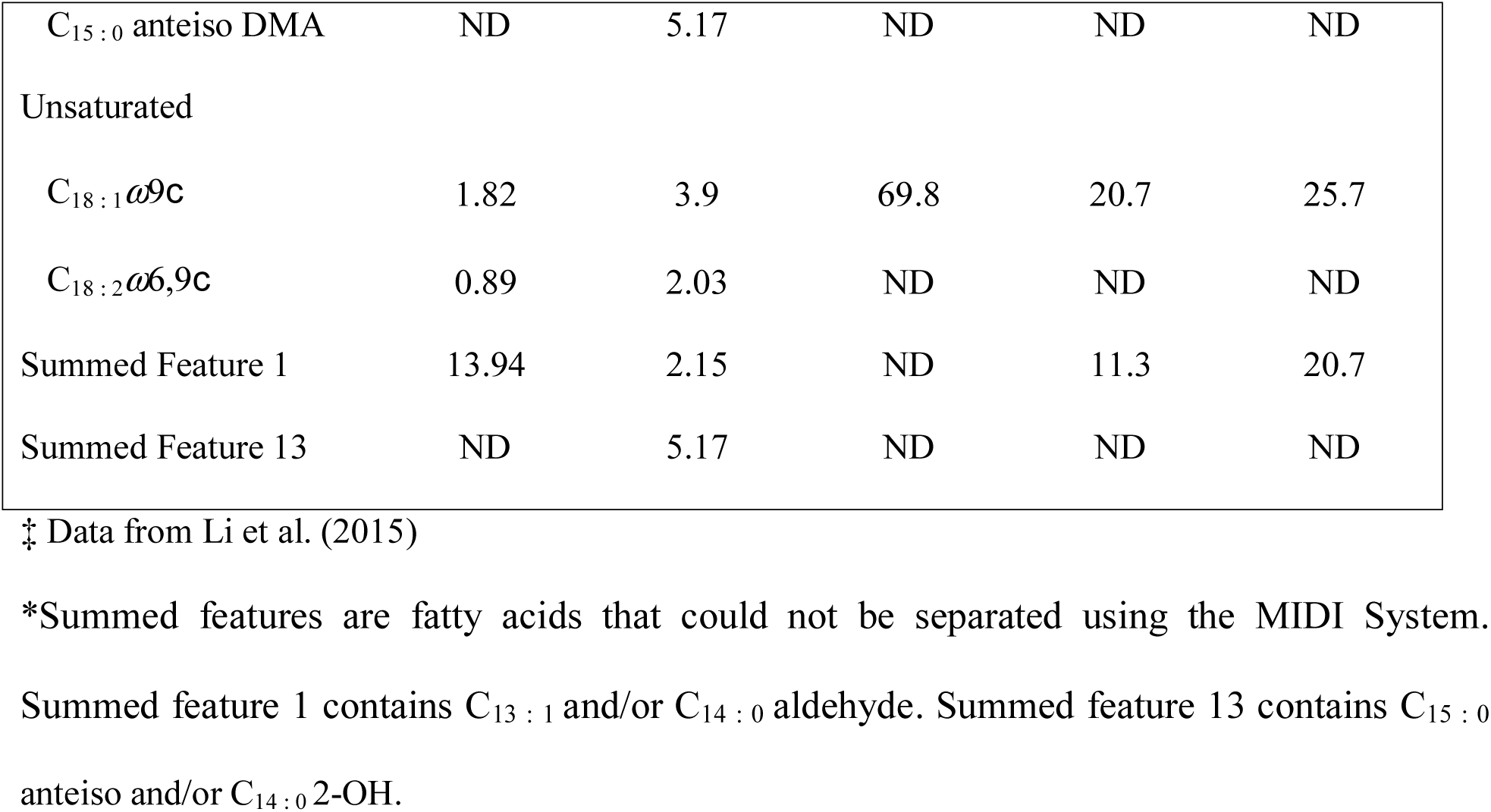
Cellular fatty acid compositions of strains SW165 and related strains of *Olsenella* Strains: 1 – SW165; 2 – *O. profusa* DSM 13989^T^; 3 – *O. uli* DSM 7084^T^; 4- *O. umbanata* DSM 22620^T^; 5- *O. scatoligenes* DSM 28304^T^. Values are percentages of total fatty acids detected. Fatty acids with contents of less than 1% in all strains are not shown; ND, Not detected.

## 16S RNA phylogeny

Genomic DNA of the strain SW165 was extracted using a DNeasy Blood & Tissue kit (Qiagen), according to the manufacturer’s instructions. The 16S rRNA gene sequences were amplified using universal primer set 27F (5’-AGAGTTTGATCMTGGCTCAG-3’; Lane et al., 1991) and 1492R (5’-ACCTTGTTACGACTT-3’; Stackebrandt et al., 1993) (18, 19), and sequenced using a Sanger DNA sequencer (ABI 3730XL; Applied Biosystems). The 16S rRNA gene sequence of SW165 was then compared with closely related strains from the GenBank (www.ncbi.nlm.nih.gov/genbank/) and EZtaxon databases (www.ezbiocloud.net/eztaxon) (20). Alignment and phylogenetic analysis were conducted using MEGA7 software (21). Multiple sequence alignments were generated using the CLUSTAL-W. Reconstruction of phylogenetic trees was carried out using the maximum-likelihood (ML) (22), maximum-parsimony (MP) (23), and neighbor-joining (NJ) (24) methods. The distance matrices were generated according to Kimura’s two-parameter model. Bootstrap resampling analysis of 1000 replicates was performed to estimate the confidence of tree topologies. Based on the results of 16S rRNA sequencing, the closest relatives of SW165 were *Olsenella profura* DSM 13989^T^ (96.33□% similarity) and *Olsenella umbonate* DSM 26220^T^ (96.18%), followed by *Olsenella uli* DSM 7084^T^ (96.03%), *Lancefieldella parvula* DSM 20469^T^ (94.29%), *Olsenella scatoligenes* DSM 28304^T^ (94.15%) and *Lancefieldella rimae* ATCC 49626^T^ (93.02%). (Figure 2).

**Fig. 2.**
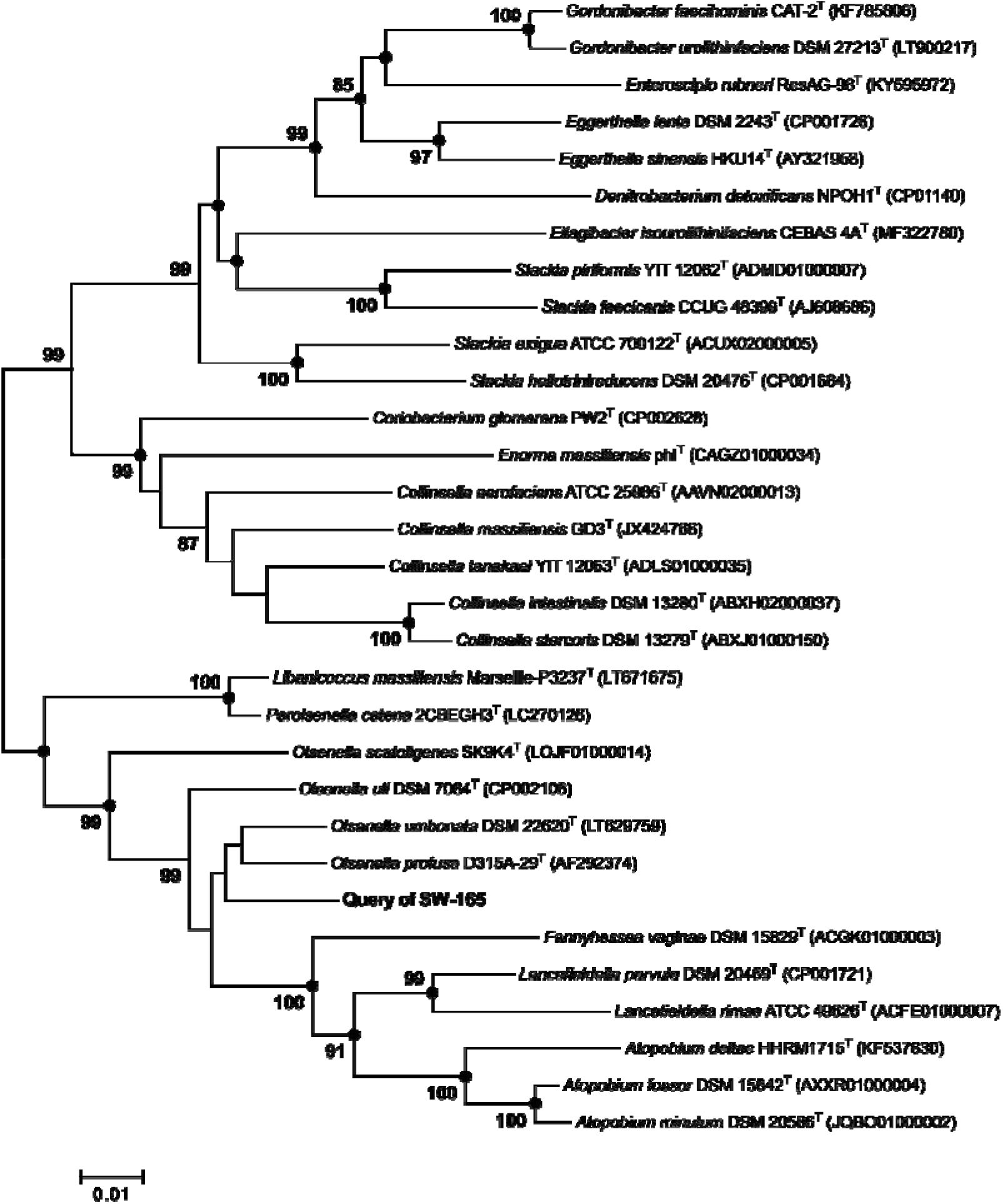
Neighbor-joining tree based on 16S rRNA gene sequences, showing the phylogenetic position of *O. lakotia* DSM 107283^T^ within closely related taxa in the genus *Olsenella* and other genera of the family *Atopobiaceae*. GenBank accession numbers of the 16S rRNA gene sequences are given in parentheses. Black circles indicate that the corresponding branches were also recovered both by maximum-likelihood and maximum parsimony methods. Bootstrap values (based on 1000 replications) greater than or equal to 70 % are shown as percentages at each node. Bar, 0.01 substitutions per nucleotide position.

## Genome Features

The whole genome sequencing of strain SW165 was performed using Illumina MiSeq sequencer using 2x 300 paired-end V3 chemistry. The reads were assembled using Unicycler that builds an initial assembly graph from short reads using the de novo assembler SPAdes 3.11.1 (25). The quality assessment for the assemblies was performed using QUAST (26). Genome annotation was performed by using Rapid Annotation using Subsystem Technology (RAST) server (27). The draft genome of strain SW165 has a total length of 2.43 Mbp with 2,228 coding sequences. The G+C content of SW165 was 67.59□mol%. Genome features of the strain were distinct from other *Olsenellae* members (Table 3). Another method of species delineation, average nucleotide identity (ANI), also confirmed that strains SW165 and DSM 13989^T^ belong to different species. The ANI of these strains was 74.18□% which was significantly less than the proposed ANI cut-off for a novel species of 95–96□% (28).

**Table 3.**
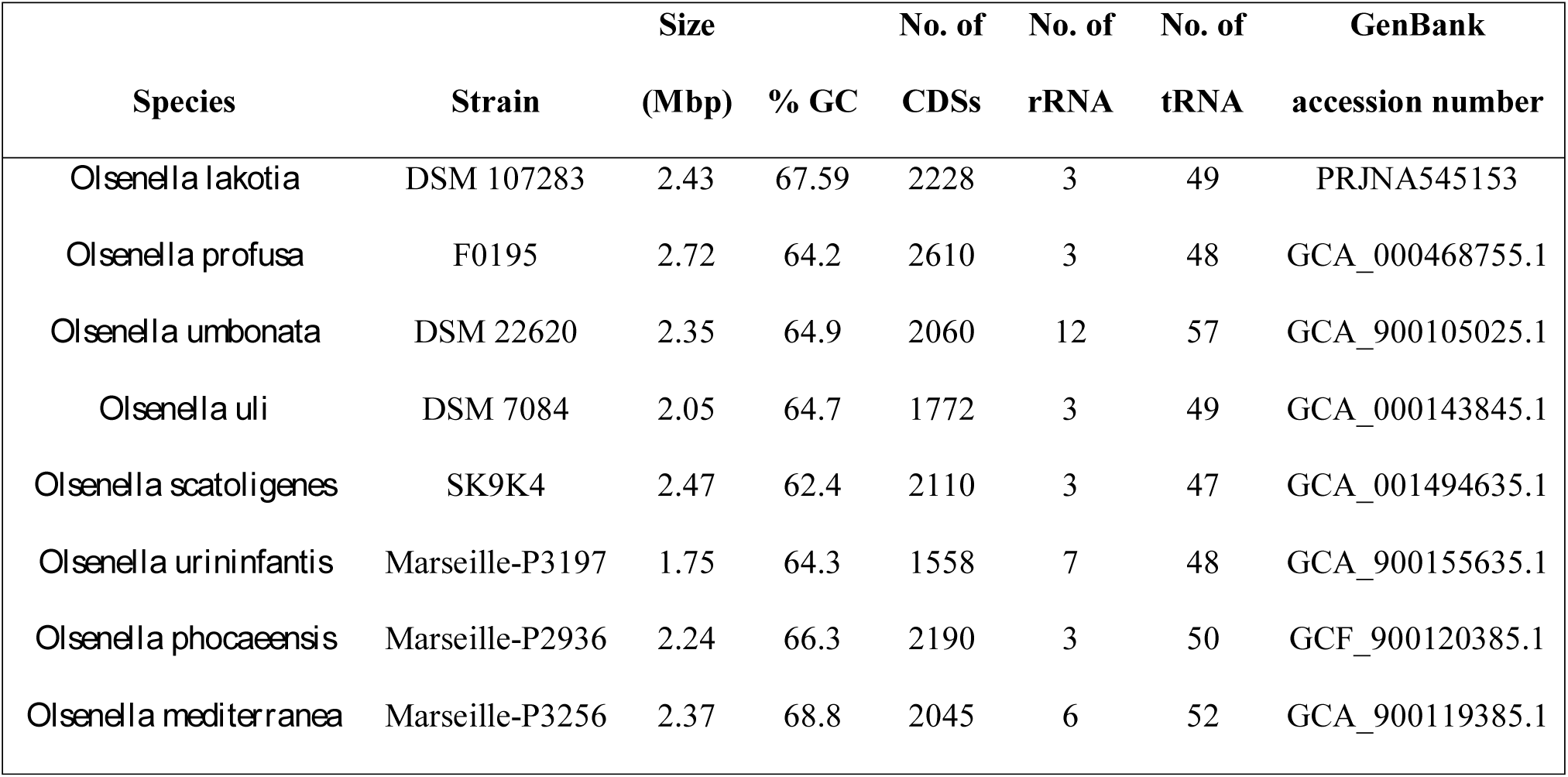
Genome characteristic of strain SW165 and other *Olsenellae*

The genotypic and phenotypic features of strains SW165 showed that this microorganism should be described as a novel species of genus *Olsenella*, for which the name *Olsenella lakotia* sp. nov. is proposed.

## Description of Olsenella SW165 sp. nov

*O. lakotia* sp. nov. (la.ko’tia N.L n. referring to native American tribe). Cells are strictly anaerobic, gram-strain-positive steptobacillus and non-motile. The average size of each cell is 0.5–2.0 µm. Colonies are visible on BHI-M agar after 2 days and are approximately 0.2–0.5 cm in diameter, cream-white, smooth, slightly umbonate with entire circular margin. The microorganism exhibits optimal growth in BHI-M medium at 45° C and pH 7. The strain utilizes arbutin, cellobiose, dextrin, D-fructode, L-fucose, D-galactose, α-D-glucose, maltose, D-mannose, D-melibiose, D-raffinose, salicin, sucrose and turanose as a carbon source. Positive enzymatic reactions are obtained for alkaline phosphatase, leucine arylamidase, cysteine arylamidase, α-galactosidase, β-galactosidase, β-glucuronidase, α-glucosidase and β-glucosidase. The volatile fatty acid produced by this strain is acetic acid. The primary cellular fatty acids are C_12□:□0_, C_14□:□0_, C_14□:□0_ DMA and summed feature 1.

The type strain SW165 (DSM 107283^T^) was isolated from the cecum of feral chicken. The DNA G+C content of the strain is 67.59 mol%.

## Prologue

The GenBank accession number for the 16S rRNA gene sequence of strain SW165^T^ is MK963074. The GenBank Biopeoject ID number for the draft genome sequence of strain SW165^T^ is PRJNA545153.

## AUTHOR STATEMENTS

### Acknowledgements

The authors would like to thank Electron Microscopy Core Facility at the Bowling Green State University, Ohio, USA for assistance with scanning electron microscopy.

### ABBREVIATIONS

ANI: average nucleotide identity

